# Membrane-Interactive Compounds from *Pistacia lentiscus* L. Thwart *Pseudomonas aeruginosa* Virulence

**DOI:** 10.1101/2020.03.17.995043

**Authors:** Ali Tahrioui, Sergio Ortiz, Onyedikachi Cecil Azuama, Emeline Bouffartigues, Nabiha Benalia, Damien Tortuel, Olivier Maillot, Smain Chemat, Marina Kritsanida, Marc Feuilloley, Nicole Orange, Sylvie Michel, Olivier Lesouhaitier, Pierre Cornelis, Raphaël Grougnet, Sabrina Boutefnouchet, Sylvie Chevalier

**Affiliations:** Université de Rouen Normandie, Normandie Université, Laboratoire de Microbiologie Signaux et Microenvironnement, LMSM EA4312, Évreux, France; Université Paris Descartes, Faculté des Sciences Pharmaceutiques et Biologiques, Équipe Produits Naturels, Analyses et Synthèses (PNAS), CiTCoM UMR 8038 CNRS, Paris, France; Centre de Recherche Scientifique et Technique en Analyses Physico-Chimiques, CRAPC, Bou Ismaïl, Algérie

**Keywords:** *Pistacia lentiscus*, Fruit-derived extract, Ginkgolic acids, Anti-virulence, *Pseudomonas aeruginosa*, Membrane stiffness, ECFσ SigX

## Abstract

*Pseudomonas aeruginosa* is capable to deploy a collection of virulence factors that are not only essential for host infection and persistence, but also to escape from the host immune system and to become more resistant to drug therapies. Thus, developing anti-virulence agents that may directly counteract with specific virulence factors or disturb higher regulatory pathways controlling the production of virulence armories are urgently needed. In this regard, this study reports that *Pistacia lentiscus* L. fruit cyclohexane extract (PLFE1) thwarts *P. aeruginosa* virulence by targeting mainly the pyocyanin pigment production by interfering with 4-hydroxy-2-alkylquinolines molecules production. Importantly, the anti-virulence activity of PLFE1 appears to be associated with membrane homeostasis alteration through the modulation of SigX, an extracytoplasmic function sigma factor involved in cell wall stress response. A thorough chemical analysis of PLFE1 allowed us to identify the ginkgolic acid (C17:1) and hydroginkgolic acid (C15:0) as the main bioactive membrane-interactive compounds responsible for the observed increased membrane stiffness and anti-virulence activity against *P. aeruginosa*. This study delivers a promising perspective for the potential future use of PLFE1 or ginkgolic acid molecules as an adjuvant therapy to fight against *P. aeruginosa* infections.

## INTRODUCTION

Bacterial infections still constitute a serious public health threat even though their prevention and treatment have been improved over the last decades. The effects of common antibiotics are no longer effective against microbial threats including *Enterococcus faecium*, *Staphylococcus aureus*, *Acinetobacter baumannii*, *Pseudomonas aeruginosa*, and *Enterobacter* species, also known as the “ESKAPE” pathogens group.^1^ In a recent report published by the World Health Organization (WHO), *P. aeruginosa* was categorized as one of the “critical priority pathogens” for which there is an urgent need for the discovery of alternative and innovative new therapies.^2^

*P. aeruginosa* is predominantly responsible for different life-threatening infections in humans, including the respiratory system, burn and wound, urinary tract as well as medical implant devices.^3^ This notorious multidrug resistant opportunistic Gram-negative bacterium deploys a wide variety of virulence factors and host-degrading enzymes as well as multiple secondary metabolites.^4^ Pyocyanin is an important virulence factor produced and secreted abundantly by nearly 95% of *P. aeruginosa* isolates.^5^ This phenazine-derived pigment, blue-green in color, confers a greenish hue to the sputum of cystic fibrosis (CF) individuals suffering *P. aeruginosa* chronic lung infection.^6^ Moreover, pyocyanin is a highly diffusible redox-active secondary metabolite which plays an important role in several physiological processes^6,7^ making it a good target. Therefore, pyocyanin production hindrance may have consequences regarding the cytotoxic effects and the full virulence of *P. aeruginosa* during infections related to airways in CF.

The sophisticated quorum sensing (QS) circuitry in *P. aeruginosa* strongly controls the biosynthesis of pyocyanin. This process starts with the synthesis of the *N*-acyl-L-homoserine lactone (AHL) type signal molecules followed by the *Pseudomonas* quinolone signaling (PQS). Next, PQS regulates the expression of *phzA-G* operons resulting in the production of phenazine-1-carboxylic acid (PCA) which is then modified to produce predominantly pyocyanin *via* the action of the enzymes encoded by *phzM and phzS*.^8^ In addition, pyocyanin biosynthesis and regulation have been also linked to SigX, an extracytoplasmic function sigma factor (ECFσ) that plays an essential role in the cell wall stress response network.^9–12^ In *P. aeruginosa*, the ECFσ SigX is a global regulator that modulates the expression of more than 300 genes, including several genes involved in pyocyanin biosynthetic pathways.^9,11,13^ More importantly, SigX has been reported to be involved in the regulation of fatty acid biosynthesis, which triggers changes in cell membrane phospholipid composition that affect membrane fluidity and envelope integrity.^11,14–16^ SigX is also involved in physiological processes as important as iron uptake, virulence, motility, attachment, biofilm formation, and antibiotic resistance and susceptibility *via* direct and/or indirect governance.^9–11,17^

In view of this, developing anti-virulence agents that may precisely hinder the production, secretion, or function of virulence determinants or interfere with their regulation has emerged as an alternative promising strategy to fight against bacterial pathogens.^18,19^ Plants have long been used in traditional medicine to prevent or treat infectious diseases in many countries.^20^ Hydrophilic and lipophilic extracts from plants were reported to contain abundant and diverse range of bioactive compounds with anti-virulence properties.^19,21,22^ *Pistacia lentiscus* L., a plant commonly known as mastic tree or lentisc, is an evergreen shrub of the family of *Anacardiaceae* widespread all around the Mediterranean area where it grows wild in a variety of ecosystems.^23,24^ Medicinal uses of the fruit, galls, resin, and leaves of *P. lentiscus* L. Are described since antiquity. However, they may differ either in the therapeutic indication or in the plant part used for medicinal purpose, depending of the geographical area.^25,26^

Currently, there are no research, which have explored the anti-virulence potential of *P. lentiscus* L. fruit, or the molecular mechanism of action, which underpins its major bioactive compounds against *P. aeruginosa*. Herein, we report that *P. lentiscus* L. fruit cyclohexane extract (PLFE1) can function as a potent pyocyanin inhibitor while being devoid of any antibacterial activity as judged by cell growth and viability assays. We show that PLFE1 interferes with 4-hydroxy-2-alkylquinolines molecules production, which might explain its anti-pyocyanin activity. We also demonstrate that PLFE1 is able to increase significantly membrane stiffness in *P. aeruginosa*. Interestingly, the cell wall stress response ECFσ SigX is found to be a key regulatory element in this phenomenon, which might respond to the presence of envelope-interactive compounds in PLFE1. Furthermore, comprehensive chemical analyses of PLFE1 and its derived fractions allowed us to identify the ginkgolic acid (C17:1) and hydroginkgolic acid (C15:0) as the major bioactive compounds and we confirmed that they are responsible of the main anti-virulence activity against the pathogen *P. aeruginosa*.

## RESULTS

### Pyocyanin production inhibition by *P. lentiscus* L. fruit extracts

Planktonic cultures of wild-type *P. aeruginosa* H103 were exposed to cyclohexane, ethyl acetate, methanol, and water *P. lentiscus* L. fruit extracts at concentrations of 100, 50, 25, 12.5, 6.25, 3.12, 1.6 and 0.8 μg mL^−1^ and assayed for interference with pyocyanin production (Fig. 1A). Pyocyanin levels from H103 strain cultures untreated with *P. lentiscus* L. fruit extracts and treated with 1% v/v DMSO were also measured as negative control and defined as 100% pyocyanin production. The PLFE1 (cyclohexane extract) showed statistically significant inhibition of pyocyanin biosynthesis at the all concentration tested. The PLFE1 at 100 μg mL^−1^ concentration caused the maximum percent reduction in pyocyanin production (82%; *P*<0.0001) over the untreated control with an IC_50_ value of 4.9 μg mL^−1^. In addition, PLFE1 inhibitory activity was shown to be dose-dependent. The PLFE2 (ethyl acetate extract) was also capable of inhibiting pyocyanin biosynthesis following the same pattern as observed for PLFE1 but not in a drastic manner where approximately 50% of inhibition of this pigment production was observed at 100 μg mL^−1^ (*P*<0.001). There was no significant or slight variation in pyocyanin production by H103 strain when exposed to PLFE3 (methanol extract) or PLFE4 (water extract). Of all the *P. lentiscus* L. fruit extracts tested for pyocyanin inhibition, PLFE1 cyclohexane extract showed remarkably better inhibitory activity and was therefore selected for all further sets of experiments. Further, the inhibitory activity of PLFE1 on phenazine biosynthetic pathway was investigated at the level of gene expression. In the presence of PLFE1 at 100 μg mL^−1^, RT-qPCR analyses results showed significant down-regulation of the *phzA* gene involved in the production of phenazine-1-carboxylic acid (PCA), *phzM*, and *phzS* which converts PCA into the derivative pyocyanin (Fig. 1B). However, the expression levels of the *phzH* gene were not significantly different in H103 strain treated with PLFE1 as compared to untreated H103 (Fig. 1B). In summary, the repression of the expression levels of genes involved in phenazine biosynthesis correlate with pyocyanin production inhibition indicating that PLFE1 has the ability to function as a strong inhibitor of the pyocyanin virulence factor in *P. aeruginosa* H103 strain.

**Figure 1.**
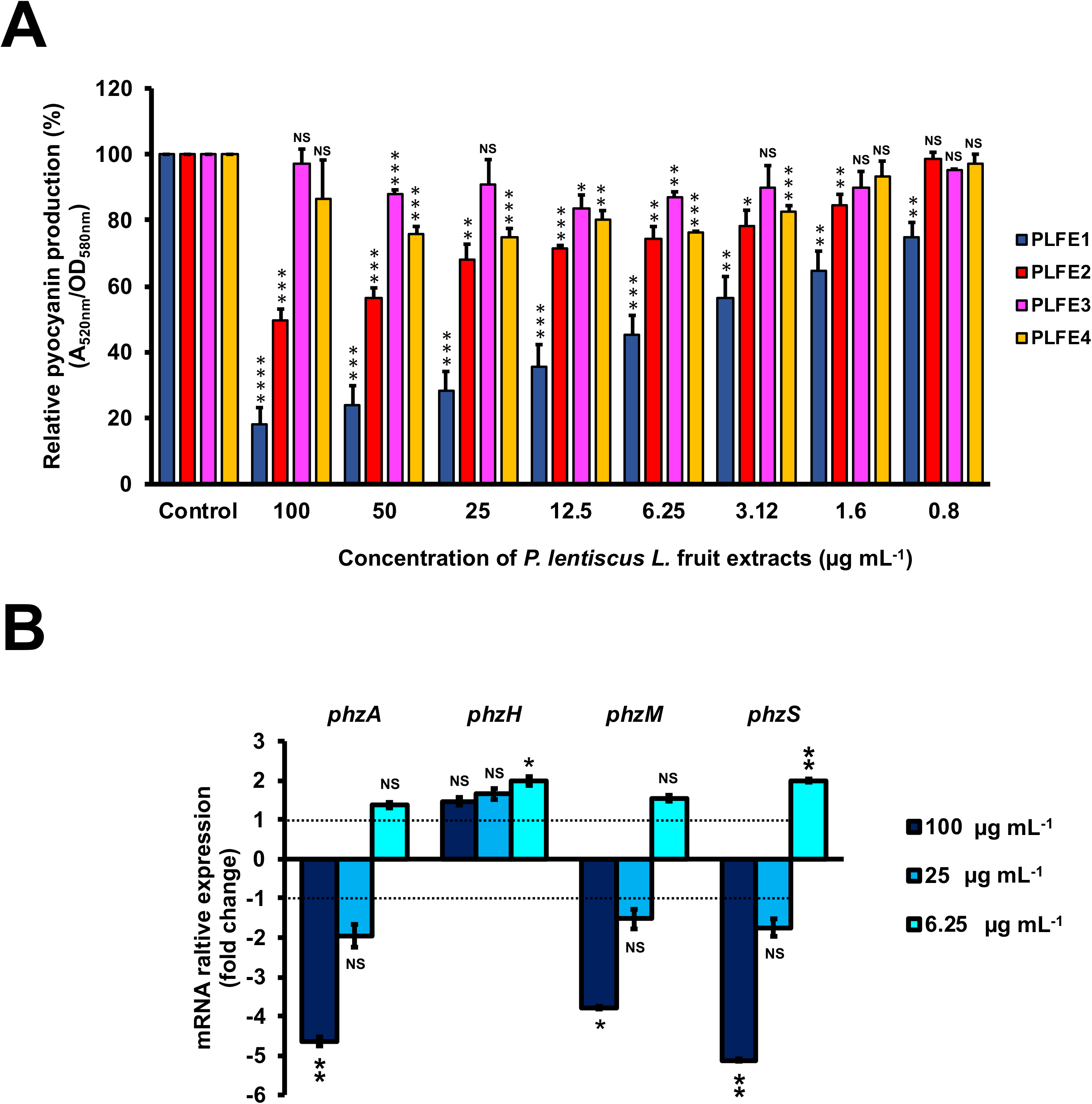
*P. lentiscus* L. fruit extract (PLFE) is a potent inhibitor of pyocyanin production in *P. aeruginosa* strain H103. (**A**) Effect of various concentrations of *P. lentiscus* cyclohexane (PLFE1), ethyl acetate (PLFE2), methanol (PLFE3), and water (PLFE4) extracts on pyocyanin production. (**B**) Relative expression levels of representative genes from phenazine biosynthesis pathway in H103 treated with PLFE1 at 100, 25, and 6.25 μg mL^−1^ compared to the relative mRNA levels in the control condition (H103 untreated). Values represent the mean (± SEM) of three independent assays. Statistics were achieved by a two-tailed *t* test: ****, *P*<0.0001; ***, *P*=0.0001 to 0.001; **, *P*=0.001 to 0.01; *, *P*=0.01 to 0.05; ^NS (Not Significant)^, *P*≥0.05.

### *P. aeruginosa* virulence attenuation by PLFE1

We next sought to evaluate anti-virulence effect of PLFE1 on *P. aeruginosa* strain H103 using human lung A549 cells and *Caenorhabditis elegans* infection models. As depicted in Fig. 2A, the results showed lower LDH release (20%; *P*≤0.05) in A549 cells after their infection with H103 treated with PLFE1 (100 μg mL^−1^) for 20h incubation. A549 cells infected with non-treated strain H103 were used as a control. This result revealed that PLFE1 exhibited significant *P. aeruginosa* anti-virulence effect in human lung A549 cells. In addition, the impact of PLFE1 on *in vivo* virulence of *P. aeruginosa* strain H103 was assessed using a *C. elegans* fast-killing infection assay. When *C. elegans* worms were placed on a lawn of H103 strain, the percentage of nematodes survival drastically decreased (65%; *P*<0.0001) after 24h incubation as compared to *C. elegans* fed with *E. coli* OP50 (Fig. 2B). Interestingly, the presence of PLFE1 (100 μg mL^−1^) significantly protected *C. elegans* from killing by *P. aeruginosa* (27%; *P*<0.0001) as compared to *C. elegans* when applied to lawns of untreated H103 (Fig. 2B). Altogether, these data indicate that PLFE1 attenuated the virulence of *P. aeruginosa* strain H103 in human lung A549 cells and *C. elegans* infection models.

**Figure 2.**
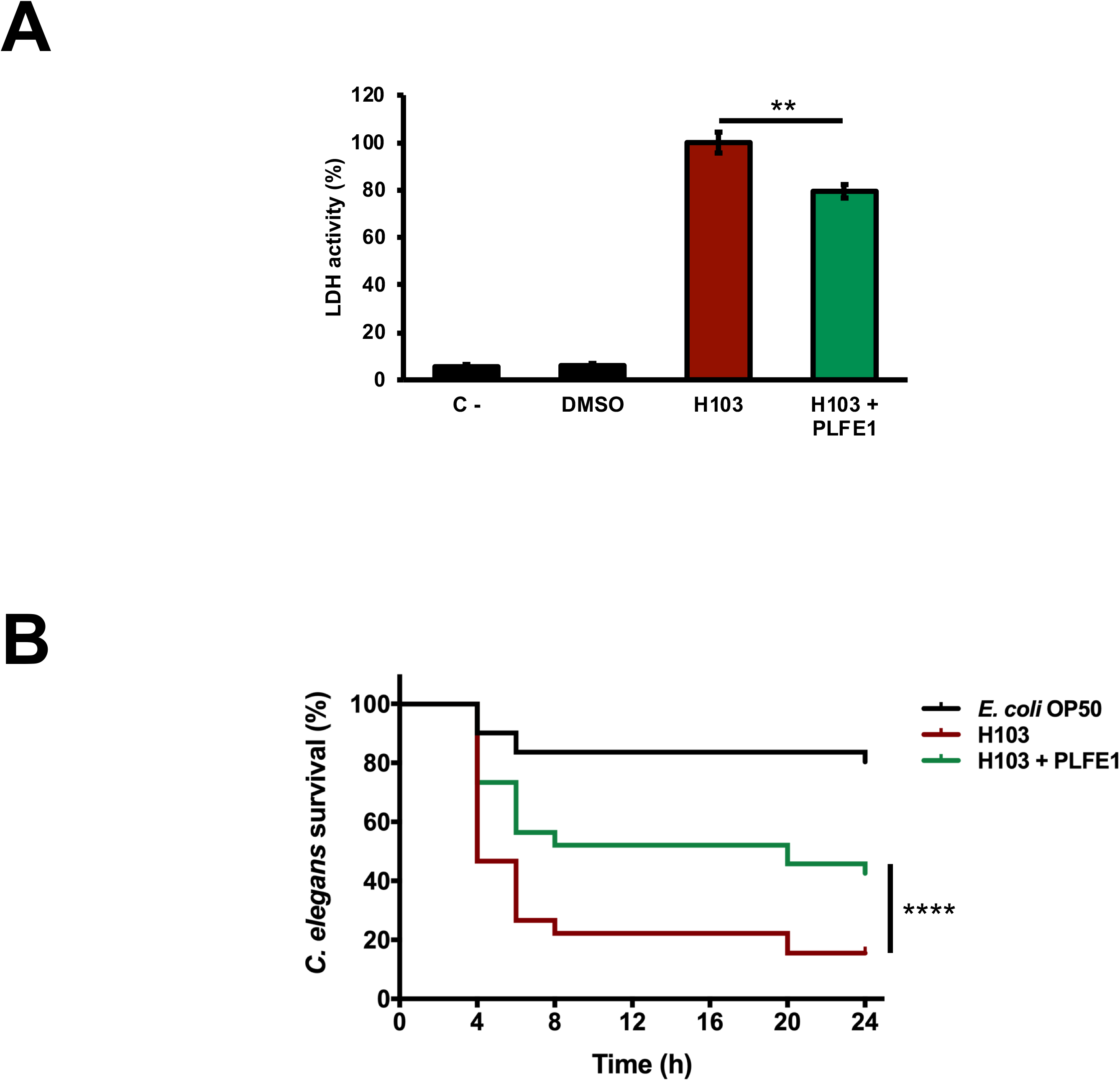
Virulence attenuation on *P. aeruginosa* by PLFE1. (**A**) Anti-virulence effects of PLFE1 in human A549 lung cells infection model. The presence of PLFE1 (100 μg mL^−1^) significantly protected A549 lung cells from lysis after 20h infection. Data are presented as the mean ± SEM values of four independent experiments performed in duplicate. ** *P*=0.001 to 0.01 (two-tailed *t* test) versus untreated cells. (**B**) *P. aeruginosa* H103 virulence attenuation in a *C. elegans* infection model by PLFE1. Sixty L4-stage nematodes per experimental group were placed on lawns of *E. coli* OP50 (black) or H103 strain in the absence (red) or presence of PLFE1 at 100 μg mL^−1^ (green). Alive nematodes were scored at 4-, 6-, 8-, 20- and 24-h after the start of the assay. ****, *P*<0.0001 (log-rank [Mantel-Cox] test) versus untreated cells.

### Absence of effect of PLFE1 on *P. aeruginosa* growth and cell viability

The impact of PLFE1 (cyclohexane extract) on the cell growth of *P. aeruginosa* H103 was monitored at 37 ºC over the course of 24 h. None of the concentrations assayed (100, 50, 25, 12.5, 6.25, 3.12, 1.6 and 0.8 μg mL^−1^) had an impact on planktonic cell growth of H103 strain as compared to the untreated culture (H103 strain grown in presence of 1 % v/v DMSO) (Supplementary Fig. S1A). Moreover, the effect of PFLE1 on H103 cell viability was evaluated by flow cytometry using *Bac*Light live/dead stain. Similarly, the results showed that at all the concentrations tested, PLFE1 did not affect the cell viability according to normalized events counting of live, injured, and dead cells when compared to the control condition (Supplementary Table S1 and Fig. S1B). Based on these results, it can be concluded that the inhibition of pyocyanin pigment production by *P. lentiscus* L. fruit extract (PLFE1) was achieved without affecting the growth of the bacteria and cell viability, at least *in vitro*.

### PLFE1 modulates the production of 4-hydroxy-2-alkylquinolines (HAQs) molecules

We next sought to assess whether the inhibition of pyocyanin production was directly due to PLFE1 on Pqs QS system that is known to tightly regulate pyocyanin biosynthesis. *P. aeruginosa* H103 cultures were exposed to different concentrations of PLFE1 (100, 50, 25, 12.5, 6.25, 3.12, 1.6 and 0.8 μg mL^−1^) and then HAQs molecules were extracted twice by ethyl acetate. The production of HAQs was determined using a PAO1 Δ*pqsA* CTX-*pqsA::lux* biosensor strain which does not produce HAQs molecules and shows response to exogenous HAQs. The H103 cultures treated with 100, 50, and 25 μg mL^−1^ of PLFE1 showed significant reduced HAQs production (Fig. 3A). Additionally, we measured the expression levels of *pqsA*, *pqsH*, *pqsL*, and *pqsR* genes representative of Pqs QS system by using RT-qPCR. The assays were performed after 24h of growth of H103 untreated or treated with 100, 25, and 6.25 μg mL^−1^ PLFE1 (Fig. 3B). The *pqsA*, *pqsH* and *pqsL* genes involved in the biosynthesis of the major molecules from the HAQs family, namely 3,4-dihydroxy-2-heptylquinoline [termed the *Pseudomonas* quinolone signal (PQS)], its precursor 4-hydroxy-2-heptylquinoline (HHQ) and 4-hydroxy-2-heptylquinoline *N*-oxide (HQNO) were all significantly repressed upon exposure to PLFE1 at 100 and 25 μg mL^−1^. However, the expression of *pqsR* (*mvfR*) encoding for the cognate receptor/regulator of HHQ and PQS molecules did not change in the presence of 100 and 25 μg mL^−1^ PLFE1. Taken together, these data indicate that PLFE1 extract represses gene expression levels of the Pqs QS system that correlates with the reduction in HAQs molecules production, suggesting that PLFE1 exhibits its anti-pyocyanin production effect possibly through an anti-QS activity.

**Figure 3.**
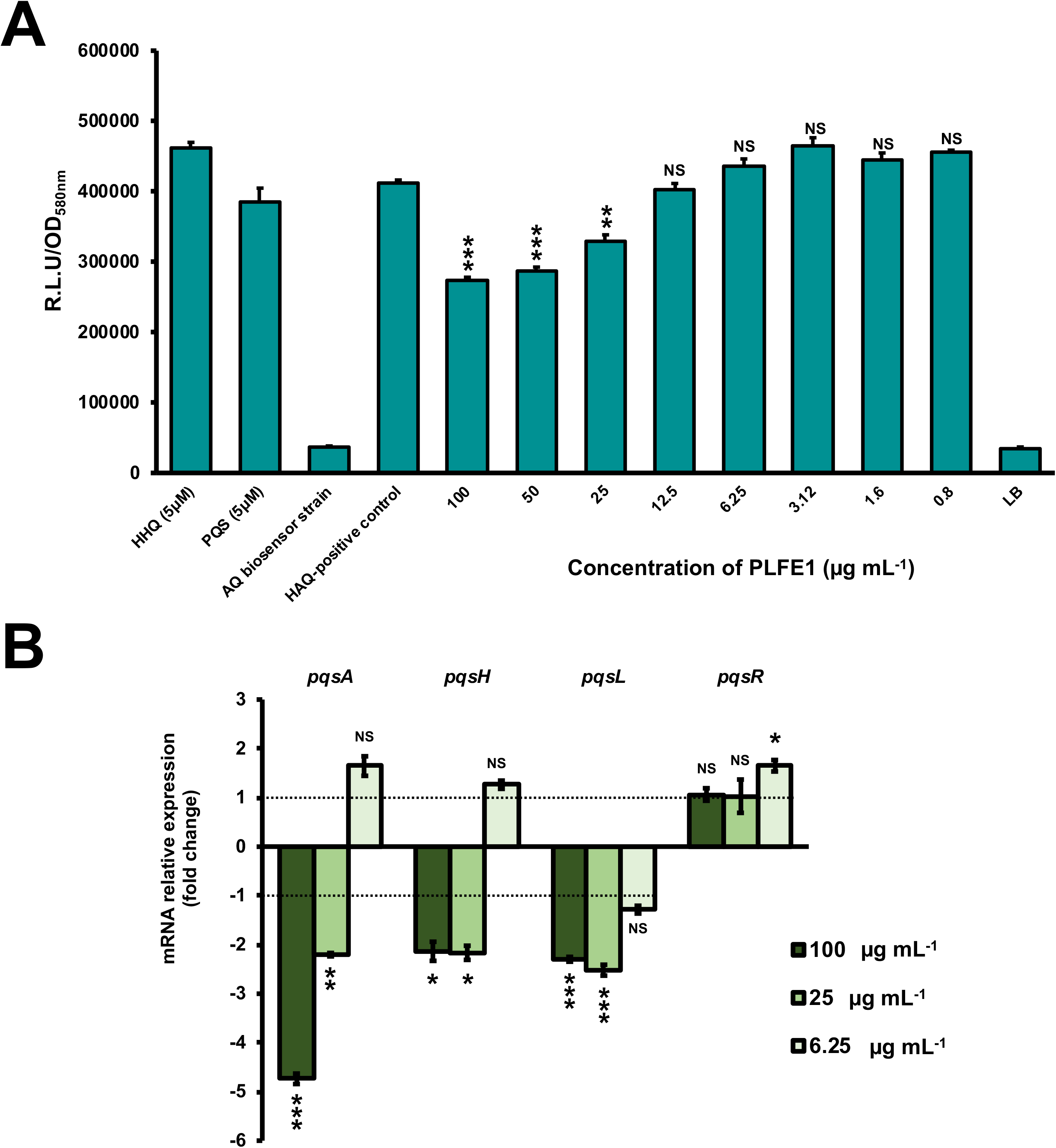
PLFE1 decreases HAQ-molecules production. (**A**) Normalized mean maximal bioluminescence output from the HAQ-biosensor strain in the presence of crude ethyl acetate HAQs extracts prepared from cultures of H103 treated with different concentrations of PLFE1 compared to positive control conditions (HHQ and PQS at 5 μM; HAQs crude ethyl acetate extract from H103 untreated) and to negative control conditions (HAQs crude ethyl acetate extract from HAQ-biosenor strain and LB medium). (**B**) Relative expression levels of representative genes from HAQ QS system in H103 treated with PLFE1 at 100, 25, and 6.25 μg mL^−1^ compared to the relative mRNA levels in the control condition (H103 untreated). Values represent the mean (± SEM) of three independent assays. Statistics were achieved by a two-tailed *t* test: ***, *P*=0.0001 to 0.001; **, *P*=0.001 to 0.01; *, *P*=0.01 to 0.05; ^NS (Not Significant)^, *P*≥0.05.

### PLFE1 leads to increased membrane stiffness

Further, we evaluated the effect of PLFE1 on *P. aeruginosa* strain H103 membrane fluidity homeostasis. Planktonic cell growths of H103 exposed to PLFE1 at concentrations of 100, 50, 25, 12.5, 6.25, 3.12, 1.6 and 0.8 μg mL^−1^ were assayed for fluorescence anisotropy (FA) using 1,6-diphenyl-1,3,5-hexatriene (DPH) fluorescent probe (Fig. 4A). Anisotropy of H103 cells increased in a concentration dependent-manner to achieve a maximum FA value of 0.223 ± 0.003 (*P*<0.001) at a concentration of 100 μg mL^−1^ of PLFE1 in comparison to the FA value of untreated control (0.186 ± 0.002). This result reflects a significant decrease of approximately 20% of membrane fluidity (membrane rigidification) revealing physiological changes and adaptations in the cellular envelope of H103 in response to PLFE1 exposure. Interestingly, mRNA expression levels of *sigX* encoding the extracytoplasmic function sigma factor (ECFσ) SigX that is required to maintain cell envelope integrity, and its known representative targets *accA*, *accB*, and *fabY* that are involved in fatty acid biosynthesis were significantly decreased upon exposure of H103 strain to PLFE1 at 100 and 25 μg mL^−1^ when compared to mRNA expression levels in the control condition (untreated H103) (Fig. 4B). Further, *sigX* expression was monitored during growth in the presence of PLFE1 at 100, 25 and 6.25 μg mL^−1^ by using a transcriptional fusion construction where the *sigX* promoter region was fused to the promoter-less *luxCDABE* cassette in the replicative pAB133 vector (pAB-P*sigX*) (Supplementary Fig. S2). Remarkably, in the presence of PLFE1, the relative bioluminescence output from H103 harbouring pAB-P*sigX* increased in a dose dependent-manner reaching significant maximal activity at 100 μg mL^−1^ during the transition from the late exponential phase to early stationary phase (Fig. 4C; Supplementary Fig. S2A). However, bioluminescence activity of pAB-P*sigX* decreased significantly in a concentration dependent-manner during the late stationary phase achieving minimal activity at 100 μg mL^−1^ of PLFE1 (Fig. 4C; Supplementary Fig. S2A), in line with the RT-qPCR results presented in Fig. 4B. These results reveal that *P. aeruginosa* exposure to PLFE1 induced increased membrane stiffness suggesting alterations in envelope homeostasis most likely through the activation of the cell wall stress ECFσ SigX. To further validate this hypothesis, we used a Δ*sigX* deletion mutant^27^ to determine the FA in the absence or presence of various concentrations of PLFE1. The FA values appeared to be unaffected in Δ*sigX* in response to PLFE1 exposure compared to the control condition (untreated Δ*sigX*) (Supplementary Fig. S2A). Altogether, these data indicate that PLFE1 contains bioactive membrane-interactive compounds that reduce membrane fluidity, in which the ECFσ SigX seems to play a key role.

**Figure 4.**
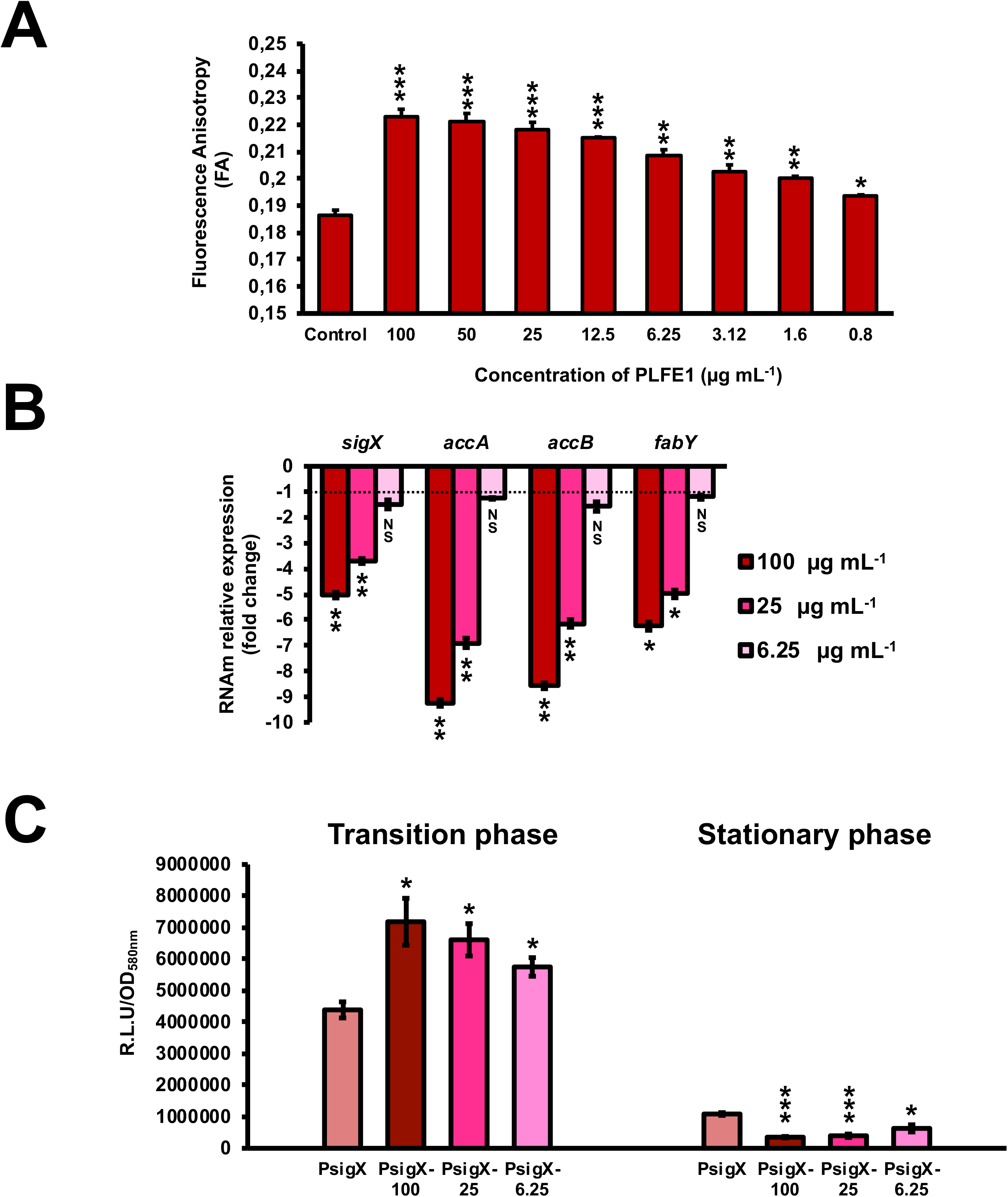
PLFE1 induces membrane stiffness. (**A**) Fluorescence anisotropy (membrane fluidity) measurements in *P. aeruginosa* strain H103 exposed to various concentrations of PLFE1 compared to the control condition (H103 untreated). (**B**) Relative expression levels of *sigX* encoding the SigX extracytoplasmic function sigma factor (ECFσ) and its known target genes (*accA*, *accB*, and *fabY*) involved in fatty acid biosynthesis in H103 treated with PLFE1 at 100, 25, and 6.25 μg mL^−1^ compared to the relative mRNA levels in the control condition (H103 untreated). (**C**) Relative bioluminescence levels of H103 harboring the pAB-P*sigX* plasmid (*sigX* promoter region) treated with PLFE1 at 100, 25, and 6.25 μg mL^−1^ compared to the relative bioluminescence levels in the control condition (H103-pAB-P*sigX* untreated). Data at both transition and stationary phases are displayed. Values represent the mean (± SEM) of three independent assays. Statistics were achieved by a two-tailed *t* test: ****, *P*<0.0001; ***, *P*=0.0001 to 0.001; **, *P*=0.001 to 0.01; *, *P*=0.01 to 0.05; ^NS (Not Significant)^, *P*≥0.05.

### Isolation and identification of major bioactive compounds of PLFE1

To isolate and identify the major constituents of PLFE1, multiple chemical analyses were performed. The fractionation of PLFE1 by medium pressure liquid chromatography (MPLC) allowed to recover a total of 13 fractions that we labelled as PLFE1-(1-13) (Supplementary Table S2). Thin layer chromatography (TLC) examination showed that the major compounds of PLFE1 were present in the fractions PLFE1-(2-4). These last were then selected for further analyses in order to identify the main compounds present in each fraction. For PLFE1-2, ^1^H-NMR spectra, proton signals between 7.36 and 6.78 ppm showed the presence of a tri-substituted aromatic ring. The signal at 5.36 ppm [t *J* = 4.6 Hz, 2H] accounted for an olefinic double bond. The major derivative ginkgolic acid (C17:1) with a molecular formula of C_24_H_38_O_3_ (*m/z* 373.1 [M-H]^−^) displayed one unsaturation in its aliphatic chain. Ozonolysis reaction of PLFE1-2 allowed to localize the double bond of ginkgolic acid (C17:1) between positions C-8’ and C-9’ (Fig. 5). LC-ESI-MS analyses corroborated the presence of ginkgolic acid (C17:1) in PLFE1-2 and showed the occurrence of hydroginkgolic acid (C15:0) derivative and a mixture of both ginkgolic acids (C17:1/C15:0) in PLFE1-3 and PLFE1-4, respectively (Fig. 5). Altogether, these results indicate that ginkgolic acid (C17:1) and hydroginkgolic acid (C15:0) are the main metabolites in PLFE1 and might be responsible for *P. aeruginosa* virulence attenuation.

**Figure 5.**
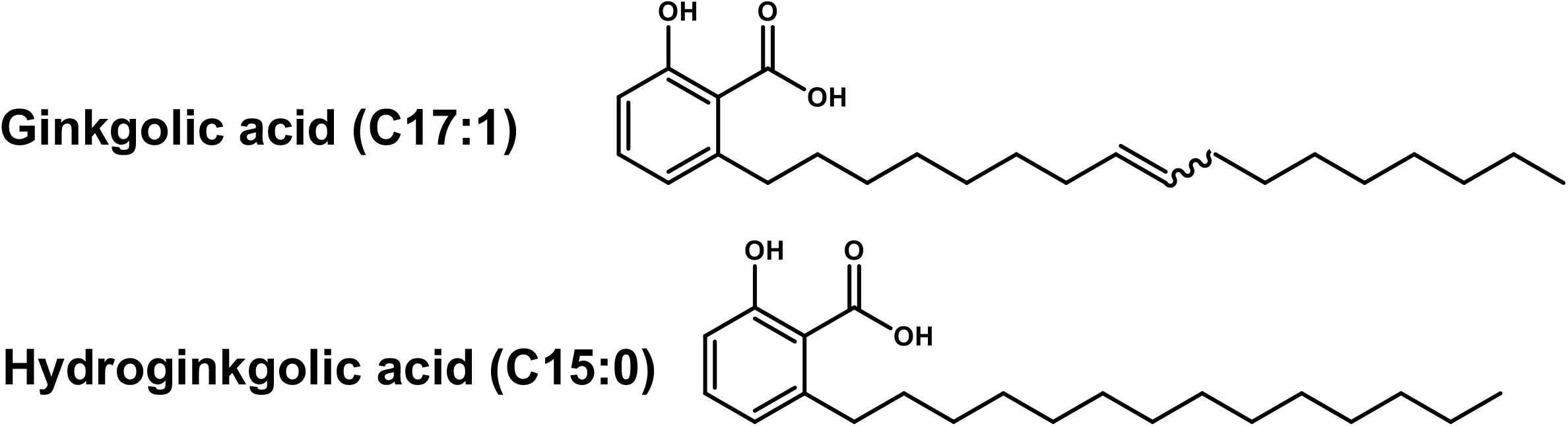
Chemical structures of ginkgolic acid derivatives identified in PLFE1-(2-4).

### Ginkgolic acid-enriched fractions from PLFE1 are involved in virulence attenuation of *P. aeruginosa*

To further assay whether GA (C17:1), GA (15:1) or a mixture of both GA (C17:1/C15:0) are the main bioactive compounds involved in the anti-virulence activity against *P. aeruginosa*, their effect on pyocyanin production and virulence attenuation in human lung A549 infection model was evaluated. Consistent with the results of pyocyanin inhibition by the crude extract PLFE1, H103 strain treated with GA-enriched fractions showed a dose-dependent reduction of pyocyanin production when compared to untreated H103 (Fig. 6A). Interestingly, the fractions enriched in GA (C17:1) or GA mix of (C17:1)/(C15:0) showed approximately 90% pyocyanin inhibition activity at 100 μg mL^−1^. The GA (C17:1) and GA (C15:0) displayed an IC_50_ of 6.3 μg mL^−1^ (16.75 μM) and 12.5 μg mL^−1^ (35.75 μM), respectively (Supplementary Fig. S3). Looking at virulence attenuation, H103 strain exposed to GA-enriched fractions showed significant reduced LDH release (20%) as compared to the control condition (H103 untreated) revealing that GA-enriched fractions were able to protect the human lung A549 line cells (Fig. 6B). Taken together, these results provide important insights into the involvement of GA-enriched fractions on the anti-virulence activity against *P. aeruginosa*.

**Figure 6.**
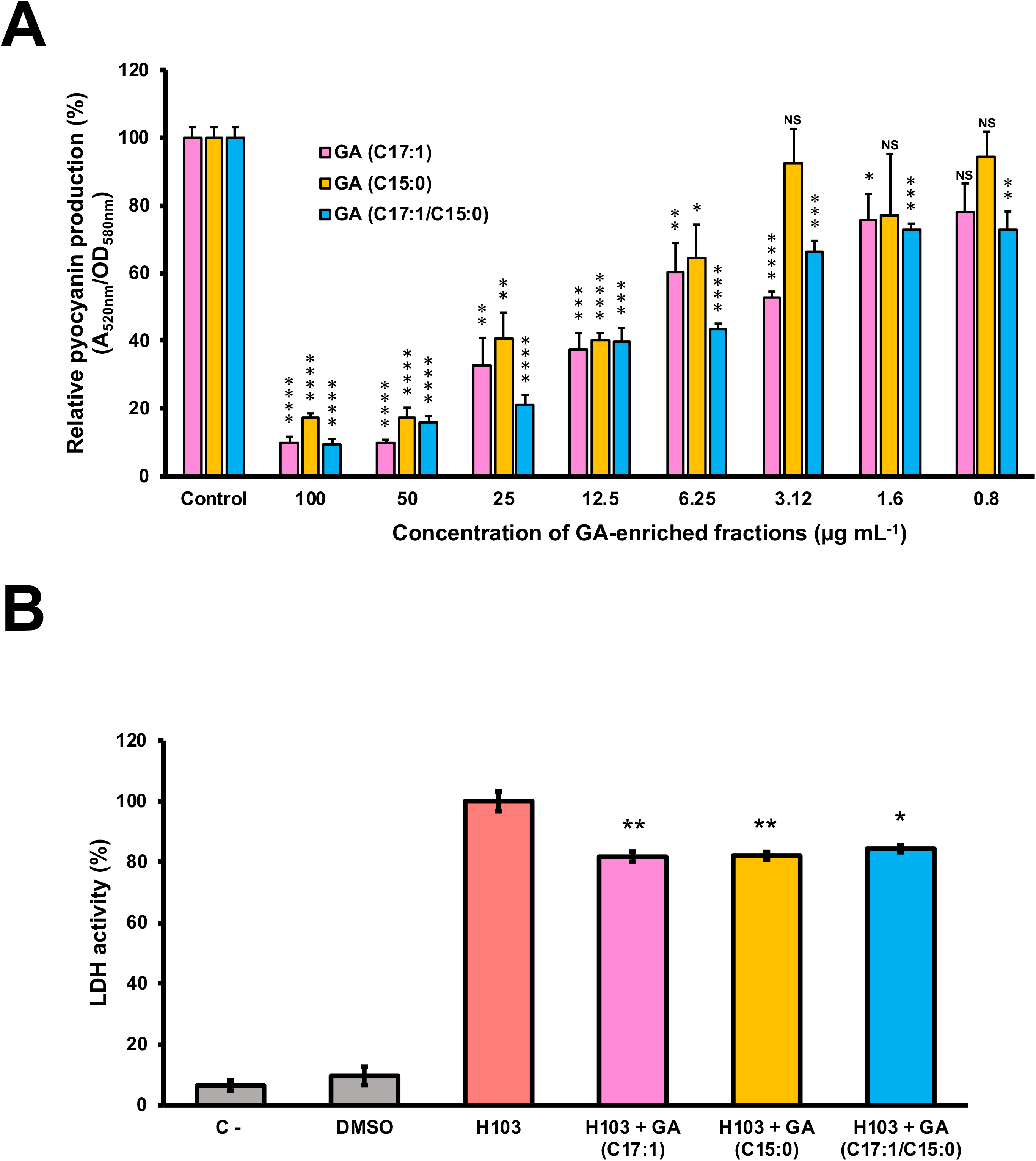
Effect of GA-enriched fractions from PLFE1 on *P. aeruginosa* virulence. (**A**) Pyocyanin production upon exposure to different concentrations of GA-enriched fractions. (**B**) Anti-virulence of GA-enriched in human A549 lung cells infection model. The presence of GA-enriched fractions (100 μg mL^−1^) significantly protected A549 lung cells from lysis after 20 h infection. Data are presented as the mean ± SEM values of four independent experiments performed in duplicate. Statistics were achieved by a two-tailed *t* test: **, *P*=0.001 to 0.01; *, *P*=0.01 to 0.05.

### Cytotoxicity of PLFE1 and GA-enriched fractions

Our last goal for this study was to investigate the potential of PLFE1 and its GA-enriched fractions to induce cytotoxicity in the human lung A549 cells. As depicted in Fig. 7A, no significant differences in cytotoxicity levels were found between A549 cells treated at various concentrations of crude extract PLFE1 after 1 h incubation as compared to the control condition (untreated A549 cells). However, a moderate cytotoxic effect was observed when A549 cells were incubated for more than 3 h in the presence of PLFE1 at concentrations above 12.5 μg mL^−1^. Remarkably, when A549 cells were treated with PLFE1 at IC_50_ value of pyocyanin activity inhibition (4.9 μg mL^−1^) there was no differences in cytotoxicity at different time points incubation assayed (1 h, 3 h, 6 h, and 24 h) (Fig. 7B). Moreover, similar interesting trend results were obtained for the GA-enriched fraction mix of both GA (C17:1/C15:0) when tested at 5.4 μg mL^−1^ IC_50_ value of pyocyanin inhibition (Fig. 7B). On the other hand, after 3 h incubation, a slight cytotoxicity increase was observed when A549 cells were exposed to enriched fractions of GA (C17:1) and GA (C15:0) separately at their IC_50_ values (6.3 μg mL^−1^ and 12.5 μg mL^−1^, respectively) (Fig. 7B). However, this cytotoxicity increased by about 3-fold after 24 h exposure as compared to the control condition (A549 cells treated with DMSO). In summary, these results indicate that PLFE1 and its complex GA-enriched fraction (C17:1/C15:0) displayed no significant cytotoxicity in the human lung A549 line cells when tested at their IC_50_ pyocyanin inhibition values.

**Figure 7.**
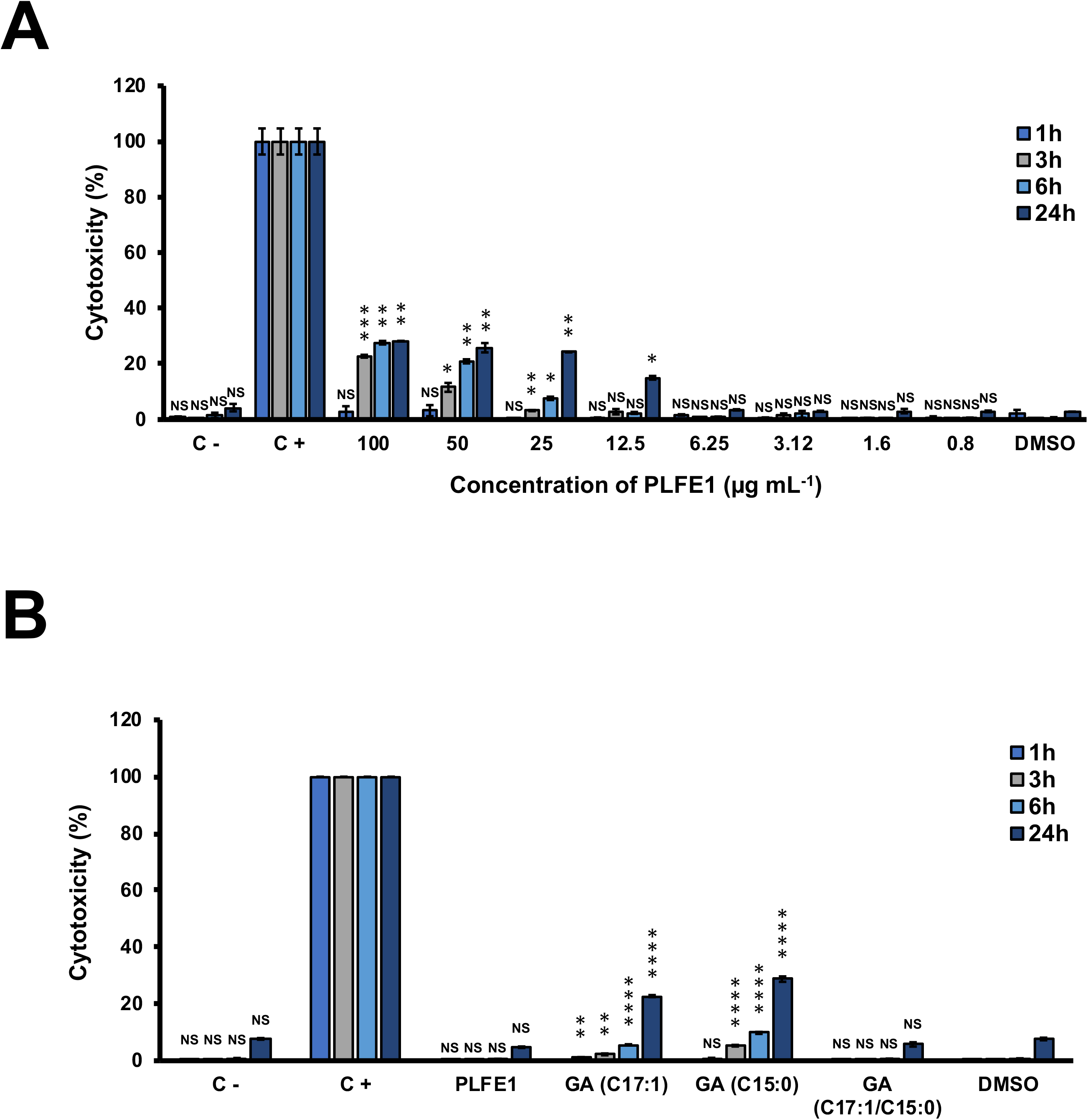
Cytotoxic effect of PLFE1 and its GA-enriched fractions on human lung A549 line cells. (**A**) PLFE1 was assayed at different concentrations. (**B**) Cytotoxicity of GA-enriched fractions exhibiting pyocyanin production inhibition. LDH release was determined at 1 h, 3 h, 6 h, and 24 h. DMSO was used as a vehicle control. Data are displayed as the mean ± SEM values of four independent experiments performed in duplicate. Statistics were achieved by a two-tailed *t* test: ***, *P*=0.0001 to 0.001; **, *P*=0.001 to 0.01; *, *P*=0.01 to 0.05; ^NS (Not Significant)^, *P*≥0.05.

## DISCUSSION

Antibiotic-resistant pathogens are threatening individuals and public health over the world to a such extent that new therapeutic strategies need to be developed urgently.^2,18^ The anti-virulence approaches represent an attracting alternative to the unmet clinical need in the treatment of bacterial infections.^28,29^ These new therapies target essentially the prevention of virulence factors production by pathogens, rather than their survival, and they also aim to thwart the regulatory mechanisms controlling their expression.^19,30^ In addition, by targeting essential virulence determinants, the emergence and spread of bacterial resistance will be reduced due to decreased selective pressure and the natural microbiota will be preserved. In line with this scenario, the current study was designed to explore new anti-virulence agents to fight against the multidrug-resistant *P. aeruginosa* that is responsible for many different infections. Thus, we based our research on the traditional medicine reported to treat several infectious diseases. In particular, the fruit of *P. lentiscus* L. is used for the treatment of respiratory tracts infections, and in ointments for articular pain, burn wounds, and ulcer among other traditional uses.^25,26^

Our findings show that *P. lentiscus* L. fruit cyclohexane extract (PLFE1) contains potent inhibitory compounds that restrict *P. aeruginosa* virulence by abolishing pyocyanin pigment production, which is known to facilitate colonization of the host and subsequent infection.^6,7^ This anti-virulence activity does not perturb growth and cell viability, which can minimize selective pressures towards the development of resistance compared to conventional antibiotics.^28^ Nonetheless, additional investigations need to be undertaken to assess whether *P. aeruginosa* can develop resistance towards PLFE1 upon repeated exposure. Further, by exploring the mechanistic behind the anti-pyocyanin activity, we have shown that PLFE1 interferes with the HAQs QS-molecules production. Since for *P. aeruginosa* infections QS is the master regulator involved in the expression of many virulence factors such as phenazines, exoproteases (elastase, alkaline protease), siderophores, and toxins among others, its inhibition represents a valuable adjuvant therapy that might be used to potentiate the activity of the available antibiotics applied to handle early *P. aeruginosa* infections.^19,30^ In the last five years there was a large amount of published investigations reporting natural products highlighting their potential in targeting bacterial virulence factors.^21^ Several plant-derived compounds that belong to alkaloids, organosulfurs, coumarins, flavonoids, phenolic acids, phenylpropanoids, terpenoids among other chemical classes have been found to be active against pyocyanin pigment production.^21^ However, there is still a significant lack on the specific underlying molecular mechanisms of action.

One important finding of the present study was that membrane stiffness in *P. aeruginosa* has significantly increased upon exposure to PLFE1. Furthermore, we demonstrate that PLFE1 decreases the expression of the ECFσ SigX, which is involved in the regulation of membrane lipid composition,^11,15^ and thus, triggers modifications on membrane fluidity and homeostasis. Accordingly, our data show a decreased expression levels of *accA* and *accB* encoding subunits of the biotin-dependent enzyme acetyl-CoA carboxylase complex (ACC) that catalyzes the first step in fatty acid biosynthesis in *P. aeruginosa*.^31^ In addition, *fabY* (PA5174) encoding the β-keto acyl synthase (FabY) involved in fatty acid biosynthesis, is another gene that showed decreased expression in presence of the PLFE1. Interestingly, these results mirror those observed in *P. aeruginosa sigX* mutant and over-expressing strains when compared to the wild-type strain.^11,15,16^ Moreover, Δ*sigX* cells display reduced membrane fluidity (membrane rigidification), HAQs QS-molecules production, pyocyanin pigment production, and virulence towards a *C. elegans* model.^9,16^ Overall, these data provide evidences that PLFE1 most likely contain membrane-interactive compounds, as it is the case for many plant extracts.^32,33^ PLFE1 interaction with membrane lipid bilayers seems to be involved in the increased membrane stiffness, which might trigger the modulation of the ECFσ SigX. Thus, our discoveries propose that PLFE1 exerts its anti-virulence activity through a new potential mechanism that at least involves the ECFσ SigX since no impact of PLFE1 was observed in membrane fluidity of the Δ*sigX* mutant strain.

Herein, the fractionation of PLFE1 led to the isolation and identification of fractions mainly enriched in C17:1 and C15:0 ginkgolic acids (GA) or a mix of both GA (C17:1/C15:0). To the best of our knowledge, the presence of these chemical constituents in *P. lentiscus* L. fruit has never been previously reported. However, according to the literature, GA has already been isolated from *Ginkgo biloba* that has long been used in traditional Chinese medicine.^34^ The GA are a mixture of several 2-hydroxy-6-alkybenzoic acid congeners that differ in carbon alkyl group length and unsaturation and are structurally similar to salicylic acid, which is reported to impact negatively the pathogenicity of *P. aeruginosa* PA14 strain by repressing pyocyanin, elastase, and protease production.^35^ A number of pharmacological effects have been attributed to GA, such as anti-bacterial, anti-fungal, insecticidal, anti-tumour, and neuroprotective effects among others.^36–39^ Moreover, GA are also proven to be effective as anti-biofilm molecules against bacterial pathogens such as *Escherichia coli* O157:H7 strain, *Staphylococcus aureus*, *Streptococcus mutans*, *Salmonella spp*., and *Listeria spp*.^40–42^ In line with our results, antibacterial activity studies show that GA did not affect Gram-negative bacteria survival, however, they exhibit strong anti-microbial activity against Gram-positive bacteria.^39,40,43,44^ Interestingly, our results indicate that PLFE1 anti-virulence properties can be mostly attributed to the identified GA-enriched fractions (C17:1, C15:0, C17:1/C15:0) since they are able to function as pyocyanin production inhibitors without negative effect on bacterial growth. Moreover, these GA-enriched fractions are capable of inducing *P. aeruginosa* membrane rigidification (*data not shown*). This result is supported by the fact that the lipid soluble components of the cell wall of Gram-negative bacteria is shown to intercept GA compounds.^43^

Following demonstration of PLFE1 and GA-enriched fractions for their *in vitro* anti-virulence properties against *P. aeruginosa*, we assayed them for *in vivo* efficacy using the human lung A549 cells and the nematode *C. elegans* as models host. In both infection models, the virulence of *P. aeruginosa* upon exposure to PLFE1 at a dose of 100 μg/mL is shown to be significantly attenuated. Further, the evaluation of the cytotoxicity of PLFE1 and GA molecules using A549 lung human cells indicated no or slight cytotoxicity at IC_50_ values. Taken together, these studies deliver a promising perspective for the potential future development of PLFE1 or GA molecules as an adjuvant agent to fight against *P. aeruginosa* infections. Nonetheless, it remains to be verified whether PLFE1 or GA molecules have an antibiotic-potentiating activity, both *in vitro* and *in vivo*, against the pathogen *P. aeruginosa*.

## CONCLUSION

Overall, the data of this study suggest that the anti-virulence efficacy of PLFE1 and GA-enriched fractions might possibly be attributed to their interaction with the lipid bilayer membranes of *P. aeruginosa* resulting in the modification of membrane fluidity. Therefore, the increased membrane stiffness in presence of PLFE1 and GA-enriched fractions appears to be mediated through the modulation of the ECFσ SigX, which is known to be involved in the regulation of *P. aeruginosa* virulence. These findings need further experimental evidences, not least in terms of identifying the specific molecular mechanisms of action leading to impact the ECFσ SigX as being a potential molecular target to alter the expression of virulence in *P. aeruginosa*.

## METHODS

### Bacterial strains, media and growth conditions

The *P. aeruginosa* H103, H103-pAB-P*sigX*, and H103-Δ*sigX*^27^ used in this study are all derivatives of *P. aeruginosa* wild-type PAO1. Planktonic cultures were grown aerobically at 37 ºC in LB broth on a rotary shaker (180 rpm) from an initial inoculum adjusted to an OD at 580 nm of 0.08. The antibiotics stock solutions used in this study were sterilized by filtration through 0.22-μm filters, aliquoted into daily-use volumes and kept at −20 ºC.

### Collection and preparation of *P. lentiscus* L. fruit extracts

Fruits of *P. lentiscus* L. were collected within the wilaya of Jijel, Algeria, in September 2016, with the collect agreement (48/MC/DGCE/DSPEC/2016) delivered by the Research Centre on Analytical Chemistry (CRAPC). A voucher herbarium specimen is deposited in the herbarium of PNAS laboratory, Université Paris Descartes (France). A sample of 8.98 g of fruits was subjected to a pressurized solvent extraction (PSE) using a Speed Extractor E-914 (Büchi) equipped with four cells (120 mL) and a collector with four flat bottom vials (220 mL) successively with cyclohexane, ethyl acetate, methanol, and water. Maximum pressure and temperature were adjusted to 100 bar and to 50 °C, respectively. Two extraction cycles with a hold-on time of 15 min were performed in each case. Solvents were removed under reduced pressure. A total of 4 extracts were obtained (Supplementary Table S3). Solutions at 10 mg mL^−1^ were prepared in DMSO of each extract and stored at 4 °C until use.

### Pyocyanin quantification assay

To perform pyocyanin pigment quantification assay, *P. aeruginosa* H103 cells untreated (grown in presence of 1% DMSO) and treated with 100, 50, 25, 12.5, 6.25, 3.12, 1.6 and 0.8 μg mL^−1^ of *P. lentiscus* L. fruit extracts (PLFE1-4) or ginkgolic acid-enriched fractions (PLFE1-(2-4)) were grown on 96-well microtiter plate for 24 h at 37 ºC on a rotary shaker (180 rpm). Then, supernatants samples were collected by centrifugation and extracted with chloroform. The chloroform layer (blue layer) was acidified by adding 0.5 M HCl. The absorbance of the HCl layer (pink layer) was recorded at 520nm using the Spark 20M multimode Microplate Reader controlled by SparkControl™ software Version 2.1 (Tecan Group Ltd.) and the data were normalized for bacterial cell density (OD_580nm_).

### Virulence attenuation of *P. aeruginosa* using human lung A549 line cells

The human lung A549 cells were cultured in Dulbecco’s Modified Eagle’s Medium (DMEM, Lonza, BioWhittaker®) supplemented with 4.5 g L^−1^ glucose, L-Glutamine, 10% heat-inactivated (30 min, 56 ºC) fetal bovine serum (FBS), and 100 Units mL^−1^ each of penicillin and streptomycin antibiotics. Cells were grown at 37 ºC under the atmosphere of 5% CO2 and 95% air with regularly medium change until a confluent monolayer was obtained. The anti-virulence effect of PLFE1 or ginkgolic acid-enriched fractions (PLFE1-(2-4)) on *P. aeruginosa* H103 was determined using an enzymatic assay (Pierce™ LDH Cytotoxicity Assay Kit, Thermo Scientific™), which measures lactate dehydrogenase (LDH) released from the cytosol of damaged A549 cells into the supernatant. After overnight incubation with *P. aeruginosa* H103 (10^8^ CFU mL^−1^) previously treated or untreated (control condition), the supernatants from confluent A549 monolayers grown on 24-well tissue culture plates were collected and the concentration of the LDH release was quantified. A549 cells exposed to 1X Lysis Buffer were used as a positive control of maximal LDH release (100% lysis) as specified by the manufacturer’s recommendations. The background level (0% LDH release) was determined with serum free culture medium. The LDH release assays were also used to determine the cytotoxicity of PLFE1 or ginkgolic acid-enriched fractions (PLFE1-(2-4)) in A549 cells upon direct exposure after 1-, 3-, 6-, and 24-h incubation.

### *Caenorhabditis elegans* fast-killing infection assay

The *C. elegans* wild-type Bristol strain N2 worms were grown at 22 °C on nematode growth medium (NGM) agar plates using *Escherichia coli* OP50 as a food source. The *P. aeruginosa*-*C. elegans* fast-kill infection assay was performed as described previously^45^ with minor modifications. Briefly, cultures of *P. aeruginosa* H103 strain untreated or treated with PLFE1 at 100 μg mL^−1^ were seeded on a 24-well plate containing in each well 1 mL of Peptone-Glucose-Sorbitol (PGS) agar. Control wells were seeded with 25 μL of *E. coli* OP50 from an overnight culture. The plate was then incubated at 37 ºC for 24 h to make bacterial lawns and then shifted to 22 ºC for 4 h. For each assay, 15 to 20 L4-synchronized worms were added to the killing and control lawns and incubated at 22 °C. Worm survival was scored at 4-, 6-, 8-, 20-, and 24-h after the start of the assay, using an Axiovert S100 optical microscope (Zeiss).

### Bacterial cell viability assays by flow cytometry

Cell viability assays of *P. aeruginosa* H103 cells untreated and treated by different concentrations of *P. lentiscus* L. fruit extract (PLFE1) were assessed by using the LIVE/DEAD™ BacLight™ Bacterial Viability and Counting Kit, for flow cytometry (Invitrogen, Molecular Probes). Briefly, *P. aeruginosa* H103 untreated and treated suspensions were diluted in filter-sterilized PBS to reach a final density of 1 × 10^6^ CFU mL^−1^, then stained following the manufacturer’s instructions. The data were acquired using CytoFlex S flow cytometer (Beckman coulter Life science). An aliquot of cells was killed with 100% ethanol and used as a control of death cells. The SYTO9 stained cells were detected by an excitation with 22 mW blue laser at 488 and with emission wavelength at 525 nm (green, with band pass filter of 40 nm), while the PI (Propidium iodide) stained cells were detected by 690 nm (red, with band pass filter of 50 nm).

### Extraction and quantification of HAQs

Planktonic cultures of *P. aeruginosa* H103 untreated (control DMSO 1%) and treated by 100, 50, 25, 12.5, 6.25, 3.12, 1.6 and 0.8 μg mL^−1^ of PLFE1 were subjected to HAQ molecules extraction following the technique described in a previous study.^46^ HAQs were quantified by a combined spectrophotometer/luminometer microplate assay using the biosensor strain PAO1 *pqsA* CTX-*lux::pqsA*.^47^ The HAQ biosensor strain was grown overnight and OD was measured and adjusted with fresh LB medium to achieve OD_580nm_ of 1. For each test well, 5 μL of HAQs crude extracts were diluted in 100 μL of LB and added to 100 μL of 1 in 50 dilution of the HAQ biosensor strain. Further, bioluminescence and OD_580nm_ were monitored in specialized white sided and clear bottom 96-well microtiter plate every 15 min for 24 h at 37 ºC using the Spark 20M multimode Microplate Reader controlled by SparkControl™ software Version 2.1 (Tecan Group Ltd.). Both HHQ and PQS synthetic standards (Sigma-Aldrich) at a final concentration of 5 μM, used as positive controls, were added to 1 in 100 dilution of the HAQ biosensor strain, as both activate bioluminescence production. The recorded bioluminescence as relative light units (R.L.U) were normalized to OD_580nm_ of cultures suspensions.

### Membrane fluidity measurement by fluorescence anisotropy

Fluorescence anisotropy analysis of *P. aeruginosa* cells were performed as previously described.^16^ Briefly, cell pellets of *P. aeruginosa* H103 untreated (control DMSO 1%) and treated by 100, 50, 25, 12.5, 6.25, 3.12, 1.6 and 0.8 μg mL^−1^ of PLFE1 were washed two times (7500 × g, 5 min, 25 °C) in 0.01 M MgSO_4_ and resuspended in the same wash solution to reach an OD_580nm_ of 0.1. One μL of a 4 mM of 1,6-diphenyl-1,3,5-hexatriene (DPH) stock solution (Sigma-Aldrich) in tetrahydrofuran was added to 1 mL aliquot of the resuspended cultures and incubated in the dark for 30 min at 37 °C to allow the probe to incorporate into the cytoplasmic membrane. Measurement of the fluorescence polarization was performed using the Spark 20M multimode Microplate Reader, equipped with an active temperature regulation system (Te-Cool™, Tecan Group Ltd.). Excitation and emission wavelengths were set to 365 nm and 425 nm, respectively, and the Fluorescence Anisotropy (FA) was calculated according to Lakowicz.^48^ Three measurements were performed for each sample and data were recorded using SparkControl™ software (Version 2.1, Tecan Group Ltd.). The relationship between fluorescence polarization and membrane fluidity is an inverse one, where increasing anisotropy values correspond to a more rigid membrane and vice versa. All values are reported as means of triplicate analyses for each experimental variable.

### Transcriptional fusion PsigX::*luxCDABE* construction and *sigX* promoter activity analysis in response to PLFE1

The promoter region of the *sigX* gene (P*sigX*) was amplified by PCR with primers PsigX-<u>*SacI*</u>-F and PsigX-<u>*SpeI*</u>-R (Supplementary Table S4,) incorporating SacI and SpeI linkers, respectively. The promoter region of *sigX* gene was fused to the *luxCDABE* cassette in the promoterless pAB133 vector.^49^ Upon amplification, DNA was digested with the appropriate restriction enzymes, and cloned into pAB133 generating pAB-P*sigX* vector. The pAB-P*sigX* plasmid was then transformed separately into one shot™ *E. coli* TOP10 competent cells (Invitrogen). The constructions were confirmed by DNA sequencing (Sanger sequencing services, Genewiz). Finally, the plasmids were transferred by electroporation into *P. aeruginosa* H103 strain. Promoter activity was analysed by monitoring bioluminescence during course time.

### Reverse transcription-quantitative PCR analyses (RT-qPCR)

Total RNAs from three independent H103 untreated and treated cultures with PLFE1 at 100, 25, and 6.25 μg mL^−1^ were isolated by the hot acid-phenol method^27^ followed by treatment with Turbo DNA-*free*™ kit (Invitrogen) according to the manufacturer’s protocol. Synthesis of cDNAs and RT-qPCR were achieved as previously described^50^ using the oligonucleotides listed in Supplementary Table S4. The mRNAs levels were calculated by comparing the threshold cycles (Ct) of target genes with those of control sample groups and the relative quantification was measured using the 2^−ΔΔCt^ method^51^ using DataAssist™ software (Applied Biosystems).

### PLFE1 fractionation

Two hundred mg of PFLE1 were subjected to fractionation by medium MPLC on 9.6 g of silice 60 M with cyclohexane/ethyl acetate as mobile phase solvent mixture with increasing polarity. A total of 13 fractions were recovered and designated PLFE1-(1-13) (Supplementary Table S2)

### Structure determination of compounds present in PLFE1-(2-4) fractions by NMR and ozonolysis

To determine the structure of the main compounds of PLFE1-(2-4), ^1^D and ^2^D NMR experiments were conducted in a Bruker 400 MHz spectrometer apparatus and recorded in deuterated chloroform (CDCl_3_). To identify the double bond position of the main compound identified in PLFE1-2, an ozonolysis reaction was carried out. Briefly, a solution of 10 mg (0.027 mmol, 1.0 equiv) of PLFE1-2 was put in a round bottom flask containing 20 mL of distilled dichloromethane and a small amount of Sudan III. A stream of ozone (O_3_) was bubbled into the solution at −78 °C until the pink solution became colorless (10 min). Then, 200 μL of dimethyl sulphide ((CH_3_)_2_S) (2.7 mmol, 100 equiv) were added to the reaction mixture and stirred for 1 h at 20 °C. From the reaction mixture, 1 μl was directly analyzed by GC-MS.

### Gas chromatography-mass spectrometry (GC-MS) analyses

The GC-MS analyses were carried out in a Hewlett Packard GC 6890/MSD 5972 apparatus equipped with HP-5 (30 m x 0.25 mm x 0.25 μm). Carried gas was Ar with a flow of 0.9 mL min^−1^, split 1:20, oven was programmed increasing from 180°C to 270°C at 8°C min^−1^ with an initial and final hold of 1 and 65 min, respectively. The inlet and GC-MS interface temperature were kept at 240 °C. The temperature of EI (electron impact) 70 eV was 220°C with full scan (80-500 m/z). The injection volume was 1 μL. Identification of constituents was achieved by comparing their mass spectra with those of the Mass Spectra Library (NIST 98) compounds.

### Statistical analyses

Statistical significance was evaluated using R (https://www.r-project.org).^52^ The data were statistically analyzed using two-sample unpaired two-tailed *t* test to calculate *P* values. The mean with standard error of the mean (SEM) of at least three independent biological experiments were calculated and plotted. *C. elegans* survival curves were prepared using R to perform a statistical log-rank (Mantel-Cox) test.

## DATA AVAILABILITY

The authors declare that all relevant data supporting the findings of the study are available in this article and its Supplementary Information files, or from the corresponding author upon request.

## ACKNOWLEDGMENTS

The LMSM is supported by the Région Normandie (France), Evreux Portes de Normandie (France) and European FEDER funds. PNAS and CRAPC are partners of the European Union’s H2020-MSCA-RISE-2015 EXANDAS (Exploitation of aromatic plants’ by-products for the development of novel cosmeceuticals and food supplements, Grant Agreement 691247). A.T. is supported by a post-doctoral fellowship from European Union (FEDER) and Région Normandie (France). D.T. is supported by a doctoral fellowship from the Région Normandie (France). C. A. was the recipient of a postgraduate fellowship from Campus France and FUNAI (Nigeria). S.O is supported by a CONICYT’s scholarship (Comisión Nacional de Investigación Científica y Tecnológica from Chile). The funders had no role in study design, data collection and interpretation, or the decision to submit this work for publication. We gratefully acknowledge Prof. Paul Williams (Centre for Biomolecular Sciences, University of Nottingham) for providing PAO1 Δ*pqsA* CTX-*pqsA*::*lux* biosensor strain and Mr. Abdelhamid Boudjerda for plant material collection.

## COMPETING INTERESTS

The authors declare no competing interests.

## AUTHOR CONTRIBUTIONS

A.T., S.C., E.B., S.B., R.G., and O.L. conceived and designed the experiments; A.T., S.O., O.C.A., N.B., O.M., and D.T. conducted experiments and analyzed the data; A.T., S.C., P.C., E.B., S.O., M.K., S.C., S.M., M.F., and N.O. contributed to the writing of the manuscript. All authors proofread the final draft and approved the final manuscript.

