## Supplementary Information for "Membrane-Interactive Compounds from *Pistacia lentiscus* L. Thwart *Pseudomonas aeruginosa* Virulence"

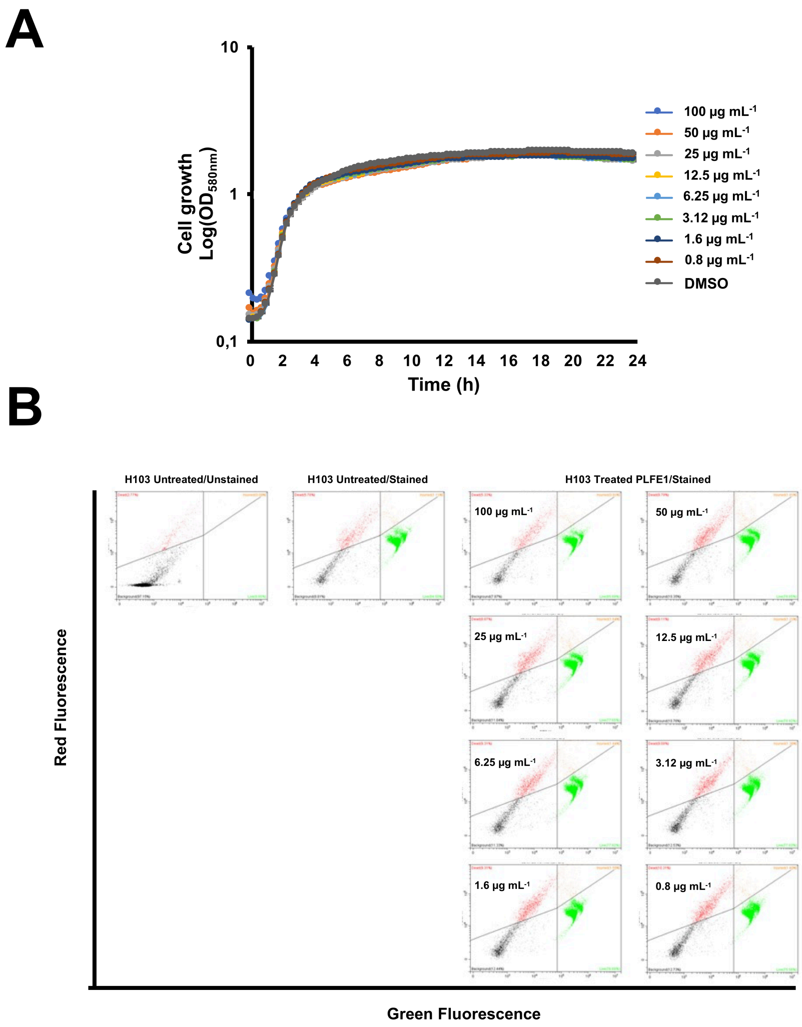


**Figure S1.** Effect of PLFE1 on *P. aeruginosa* growth and cell viability. (**A**) Growth kinetics of *P. aeruginosa* strain H103 treated with PLFE1 at various concentrations compared to the growth in the control condition (H103 untreated). Each point indicates the mean (± SEM) of $OD_{580nm}$ values. (**B**) Flow cytometry analysis of H103 cell viability upon exposure to PLFE1 at various concentrations. Suspensions at a density of $1 \times 10^6$ CFU $mL^{-1}$ were stained using the LIVE/DEAD™ BacLight™ Bacterial Viability and Counting Kit and then analyzed by flow cytometry. Live and dead cells emitting a green or a red fluorescence are indicated within a green or a red frame, respectively. The orange frame corresponds to cells with damaged membranes. The black frame corresponds to the background noise. Cytograms are representative of three independent experiments.

**Table S1.** Effect of PLFE1 on cell viability as determined by flow cytometry

| **H103 cells** | **Normalized Live Events μL$^{-1}$** | **Normalized Injuried Events μL$^{-1}$** | **Normalized Dead Events μL$^{-1}$** |
|---|---|---|---|
| Untreated | 0.963 ± 0.007 | 0.006 ± 0.001 | 0.029 ± 0.007 |
| Treated with PLFE1 (μg mL$^{-1}$) | | | |
| 100 | 0.965 ± 0.006[a] | 0.005 ± 0.001[a] | 0.029 ± 0.007[a] |
| 50 | 0.955 ± 0.003[a] | 0.008 ± 0.002[a] | 0.037 ± 0.002[a] |
| 25 | 0.958 ± 0.001[a] | 0.006 ± 0.001[a] | 0.035 ± 0.002[a] |
| 12.5 | 0.960 ± 0.002[a] | 0.005 ± 0.000[a] | 0.034 ± 0.002[a] |
| 6.25 | 0.953 ± 0.006[a] | 0.005 ± 0.001[a] | 0.042 ± 0.007[a] |
| 3.12 | 0.952 ± 0.002[a] | 0.005 ± 0.001[a] | 0.043 ± 0.002[a] |
| 1.6 | 0.951 ± 0.002[a] | 0.005 ± 0.001[a] | 0.043 ± 0.002[a] |
| 0.8 | 0.941 ± 0.006[a] | 0.006 ± 0.001[a] | 0.053 ± 0.006[a] |

[a] Not Significant ($P$≥0.05)

A

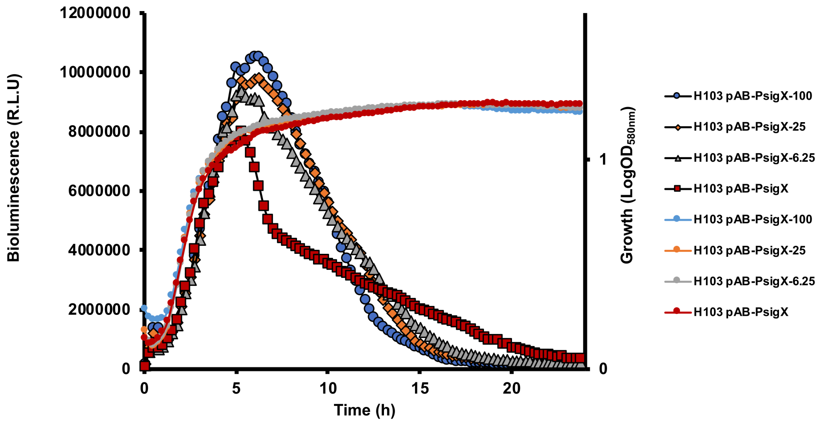


B

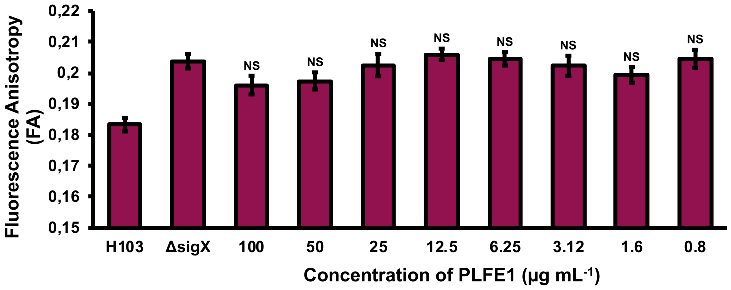


**Figure S2.** Effect of PLFE1 on the ECFσ SigX. (**A**) Relative bioluminescence levels of H103 strain harboring the pAB-P*sigX* plasmid (*sigX* promoter region) treated with PLFE1 at 100, 25, and 6.25 µg $mL^{-1}$ compared to the relative bioluminescence levels in the control condition (H103 untreated). Growth kinetics are also displayed. (**B**) Fluorescence anisotropy (membrane fluidity) measurements in *P. aeruginosa* Δ*sigX* exposed to various concentrations of PLFE1 compared to the control condition (Δ*sigX* untreated). Values represent the mean (± SEM) of three independent assays. Statistics were achieved by a two-tailed *t* test: $^{\text{NS (Not Significant)}}$, $P \geq 0.05$.

**Table S2**. Fractions of PLFE1 obtained from MPLC

| Code | Mass [mg] | Percentage [m/m, %] |
|---|---|---|
| PLFE1-1 | 3.0 | 1.5 |
| PLFE1-2 | 35.5 | 17.8 |
| PLFE1-3 | 16.5 | 8.2 |
| PLFE1-4 | 16.2 | 8.1 |
| PLFE1-5 | 16.8 | 8.4 |
| PLFE1-6 | 9.6 | 4.8 |
| PLFE1-7 | 4.2 | 2.1 |
| PLFE1-8 | 21.3 | 10.6 |
| PLFE1-9 | 5.3 | 2.6 |
| PLFE1-10 | 1.2 | 0.6 |
| PLFE1-11 | 8.7 | 4.4 |
| PLFE1-12 | 6.1 | 3.0 |
| PLFE1-13 | 2.9 | 1.4 |

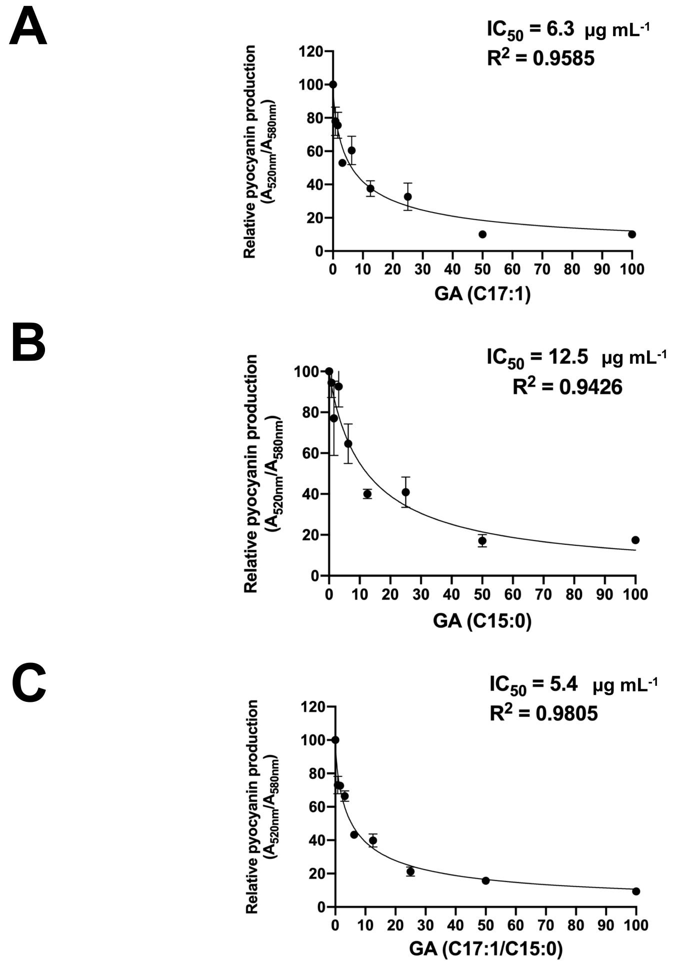


**Figure S3.** Determination of $IC_{50}$ values for GA-enriched fractions. (**A**) GA (C17:1). (**B**) GA (C15:0) (**C**) mixture of GA (C17:1/C15:0).

**Table S3.** Collection and preparation of *P. lentiscus* L. fruit extracts

| Plant material | Date of collection | Place | GPS position | Code | Solvent | Percent Yield |
|---|---|---|---|---|---|---|
| *P. lentiscus* L. fruit (PLF) | 20/09/2016 | Wilaya Jijel (Algeria) | 36° 43' 25.248'' N 5° 52' 22.4148'' E | PLFE1 | Cyclohexane | 2.7 |
| | | | | PLFE2 | Ethyl acetate | 0.6 |
| | | | | PLFE3 | Methanol | 18.2 |
| | | | | PLFE4 | Water | 8.3 |

**Table S4.** List of primers used in this study

| PA number | Gene name | Primer name | Sequence (5' > 3')[a] | Length |
|---|---|---|---|---|
| Construction of the transcriptional fusion pAB-P*sigX* | | | | |
| *PA1775-PA1776* | *cmpX-sigX* | P*sigX*-SacI-F | taataa-**GAGCTC-**gagtcgctcggcctgca | 29 |
| | | P*sigX*-SpeI-R | taataaa-**CTAGTG-**gtggaacagctccgagtgcg | 33 |
| Quantification of mRNA levels by RT-qPCR | | | | |
| *PA4210* | *phzA* | *phzA-F* | AACCACTACATCCATTCCTTCG | 22 |
| | | *phzA-R* | CGGCTATTCCCAATGCAC | 18 |
| *PA0051* | *phzH* | *phzH-F* | CGCGGGTTGGGTGGAT | 16 |
| | | *phzH-R* | ATGACCGATACGCTCGCC | 18 |
| *PA4209* | *phzM* | *phzM-F* | GCTGCGCGTAATTTGATACAAG | 22 |
| | | *phzM-R* | GATCCCGCTCTCGATCAGATC | 21 |
| *PA4217* | *phzS* | *phzS-F* | CCTGCGCGAATACGAAGAAG | 20 |
| | | *phzS-R* | CGGCCCATTCCTCTTTTTC | 19 |
| *PA0996* | *pqsA* | *pqsA-F* | CGGAGTTGCTGGCATTGC | 18 |
| | | *pqsA-R* | CTGTTGCCCATGCCATAGC | 19 |
| *PA2587* | *pqsH* | *pqsH-F* | CTCCATCGTGCAGATCCT | 18 |
| | | *pqsH-R* | CGGAATGACGCAAGGTC | 17 |
| *PA4190* | *pqsL* | *pqsL-F* | CGGTATCGCCTCCTACGTG | 19 |
| | | *pqsL-R* | GGAAGCTCACCACCAGTCG | 19 |
| *PA1003* | *pqsR* | *pqsR-F* | AACCTGGAAATCGACCTGTG | 20 |
| | | *pqsR-R* | TGAAATCGTCGAGCAGTACG | 20 |
| *PA1776* | *sigX* | *sigX-F* | AATTGATGCGGCGTTACCA | 19 |
| | | *sigX-R* | CCAGGTAGCGGGCACAGA | 18 |
| *PA3639* | *accA* | *accA-F* | TCTTCGGCAATCTGACCAGTT | 21 |
| | | *accA-R* | GTAGCCGATGTAGTCGAGGGTA | 22 |
| *PA4847* | *accB* | *accB-F* | AAGCCATGAAGATGATGAACC | 21 |
| | | *accB-R* | CGTTCTCCACCAGGATCGA | 19 |
| *PA5174* | *fabY* | *fabY-F* | AGGGCGACCTGGAGATCAT | 19 |
| | | *fabY-R* | GCGCGTCCTTCTTGTATACCA | 21 |
| | *16S* | *16S-F* | AACCTGGGAACTGCATCCAA | 20 |
| | | *16S-R* | CTTCGCCACTGGTGTTCCTT | 20 |

[a]All the primers used in this study were synthesized by Eurogentec and are based on *P. aeruginosa* PAO1 genome sequence (http://www.pseudomonas.com). Bold nucleotides indicate restriction endonuclease sites inserted within primer sequences.
